# Warm temperature inhibits cytoplasmic incompatibility induced by endosymbiotic *Rickettsiella* in a spider host

**DOI:** 10.1101/2024.04.17.589895

**Authors:** Jordyn D. Proctor, Virginija Mackevicius-Dubickaja, Yuval Gottlieb, Jennifer A. White

## Abstract

Bacterial endosymbionts manipulate reproduction in arthropods to increase prevalence in the host population. One such manipulation is cytoplasmic incompatibility (CI), wherein the bacteria sabotage sperm in infected males to reduce hatch rate when mated with uninfected females, but zygotes are “rescued” when that male mates with an infected female. In the spider *Mermessus fradeorum* (Linyphiidae), *Rickettsiella* symbionts cause variable levels of CI. We hypothesized that temperature affects CI strength and rescue in *M. fradeorum*, potentially mediated by bacterial titer. We reared *Rickettsiella*-infected spiders in two temperature conditions (26°C vs 20°C) and tested CI induction in males and CI rescue in females. In incompatible crosses between infected males and uninfected females, hatch rate from warm males was doubled (Mean±S.E. = 0.687±0.052) relative to cool males (0.348±0.046), indicating that CI induction is weaker in warm males. In rescue crosses between infected females and infected males, female rearing temperature had a marginal effect on CI rescue, but hatch rate remained high for both warm (0.960±0.023) and cool females (0.994±0.004). Bacterial titer as measured by qPCR was lower in warm than cool spiders, particularly in females, suggesting that bacterial titer may play a role in causing the temperature-mediated changes in CI.

## INTRODUCTION

Maternally-inherited bacterial symbionts are common in arthropods, some of which can manipulate host reproduction to increase bacterial prevalence within the host population. Cytoplasmic incompatibility (CI) is the most common and well-studied manipulation, sabotaging matings between infected males and uninfected females, preventing the production of uninfected offspring, and increasing the relative proportion of infected offspring in the population over time (Shropshire et al. 2020). CI is typically viewed as a modification/rescue system: The symbiont induces CI by modifying male gametes, which cannot successfully fertilize and develop into progeny unless the egg is also infected and contains an appropriate rescue factor (Shropshire et al. 2020). Numerous bacterial clades have been shown to cause CI, including *Wolbachia* (Alphaproteobacteria; reviewed in Shropshire et al. 2020), *Cardinium* (Bacteroidetes; Hunter et al. 2003), *Mesenet* (Alphaproteobacteria; Takano et al. 2021), *Spiroplasma* (Mollicutes; Pollman et al. 2022), and *Rickettsiella* (Gammaproteobacteria; Rosenwald et al. 2020). The mechanistic underpinnings of CI are only well characterized for *Wolbachia* (Shropshire et al. 2020), but genomic and phenotypic studies with other symbionts suggest novel modes of action may be present (Poinsot et al. 2003, Penz et al. 2012, Pollmann et al. 2022). Consequently, caution should be used when extrapolating inferences from *Wolbachia* CI studies to other bacterial symbionts.

There is growing evidence that symbiont-induced host phenotypes, such as CI, can be thermally sensitive but potentially idiosyncratic (Corbin et al. 2017, Doremus et al. 2019, Shropshire et al. 2020, Chrostek et al. 2021, Corbin et al. 2021, Hague et al. 2022, Jones et al. 2023, Martins et al. 2023). *Wolbachia*-induced phenotypes generally appear to weaken with increased temperature (e.g., Clancy and Hoffmann 1998, Ross et al. 2017), which likely has consequences for the global deployment of this symbiont in vector management programs (Ross et al. 2023). Other symbionts, such as *Spiroplasma*, show a reversed pattern, with weaker effects on their hosts at decreased temperature (Anbutsu et al. 2008, Corbin et al. 2021). The effect of temperature on *Rickettsiella*-induced CI has not previously been investigated, but in a non-CI-inducing *Rickettsiella* strain in aphids, temperature affects *Rickettsiella* titer, host phenotype, and spread in the host population (Gu et al. 2023). More generally, several studies have suggested that thermal conditions might limit the geographic distribution of symbionts, their phenotypes, and consequently even their host species (Zhang et al. 2019, Ross et al. 2020, Hague et al. 2022). Thus, under global climate change, it becomes critical to understand thermal effects on arthropod symbionts, because symbiont-induced phenotypes are likely to change and affect the spread and performance of both pest and beneficial arthropod hosts.

Here, we test whether temperature affects the strength of *Rickettsiella*-induced CI and bacterial titer in the linyphiid spider *Mermessus fradeorum* (Berland, 1932). Few studies of manipulative symbionts have been conducted in non-insect arthropods, which can be infected with well-known symbiotic taxa such as *Wolbachia*, and lesser known manipulative clades such as *Rickettsiella*. Populations of *Mermessus fradeorum* are composed of individuals that are naturally co-infected with different combinations of several such maternally inherited symbionts, of which only *Rickettsiella* has been shown to induce CI (Curry et al. 2015, Rosenwald et al. 2020). The mechanism by which *Rickettsiella* induces CI is unknown; however, we have observed the strength of CI to be quite variable. We hypothesized that temperature mediates the penetrance of the phenotype, potentially mediated by bacterial titer. Using laboratory-reared colonies of spiders that were either infected only with *Rickettsiella* or were uninfected because they came from lineages that had been experimentally cured via antibiotics (Rosenwald et al. 2020), we reared spiders under two temperature regimes (26°C or 20°C) and evaluated the ability of infected males to induce CI and infected females to rescue CI as a function of temperature. Because the strength of symbiont-induced phenotypes is often mediated by bacterial quantity (Lopez-Madrigal and Duarte 2019), we also evaluated bacterial titer as a function of temperature.

## EXPERIMENTAL PROCEDURES

To test the effect of temperature on *Rickettsiella*-induced CI in *M. fradeorum*, we conducted separate experiments on CI-induction in males versus CI-rescue in females. We assessed induction versus rescue separately to maximize experimental power on informative crosses. We selected two temperature treatments, warm (26°C) and cool (20°C) that fall within the range of temperatures the spiders would experience in the field, and at which we knew the spiders could be successfully reared and propagated. *Rickettsiella*-infected (R-infected) spiders were reared under both temperature regimes, while uninfected spiders were only reared under cool conditions because the experiments were designed to test for effects of temperature on CI, not on the spiders *per se*.

A cohort of R-infected (n=17) and uninfected (n=13) mother *M. fradeorum* were mated and allowed to lay eggmasses over the span of 2-3 weeks. As each clutch of spiderlings hatched, they were split into individual 4 cm diameter rearing cups with moistened plaster at the bottom for humidity control (Rosenwald et al. 2020). Half of the R-infected spiderlings were randomly assigned to be reared in a 20°C environmental chamber and half in a 26°C chamber; all uninfected spiderlings were reared at 20°C. All were maintained on a diet of *Sinella curviseta* collembola when young and *Drosophila melanogaster* fruit flies when older (Rosenwald et al. 2020). After spiders reached maturity, they were randomly assigned to mating crosses in either male CI induction or female CI rescue experiments. All experimental spiders were 7-10 weeks of age at mating; these spiders typically live 6-10 months under laboratory conditions, hence age differences of a couple of weeks are trivial.

For the male CI induction experiment, the experimental test cross was to compare CI induced by R-infected males reared under warm versus cool conditions. These males (n=30 per treatment) were mated to uninfected females to evaluate CI induction. To ensure that hatch rate reductions observed in the CI crosses were not due to innate male infertility, these same males were mated one week later to cool-reared R-infected females (compatible cross). Male *M. fradeorum* are capable of mating multiple times and we have not detected decreases in fertility or CI strength with subsequent matings (J. White, unpublished data). To control for potential differences in innate female fertility, additional sets of cool-reared R-infected (n=15) and uninfected (n=15) females were mated with uninfected males.

For the female CI rescue experiment, the experimental test cross of interest was to compare CI rescue by R-infected females reared under warm versus cool conditions. These females (n=30 per treatment) were mated to cool-reared R-infected males to evaluate CI rescue. Because we could not use the same re-mating technique with females as males, we mated separate sets of warm and cool R-infected females (n=15 per treatment) to uninfected males, as controls for female fertility. We included an additional control cross between cool R-infected males and uninfected females (n=15), to ensure that males used in this experiment were capable of causing CI. Finally, an additional set of warm-reared R-infected males (n=15) was also crossed with uninfected females to aid in comparison across the two experiments.

After matings for each experiment were complete, male spiders were placed in 1.5mL microcentrifuge tubes and stored at their rearing temperature without food for five to seven days to ensure that gut contents had been emptied. Specimens were then preserved in 95% ethanol and stored at -20°C for further analysis. Female spiders were maintained under standard rearing conditions at their assigned temperature until they deposited three eggmasses. Then, females were moved to microcentrifuge tubes and underwent the same food deprivation and preservation sequence as the males. To validate expected infection status, we extracted DNA from a subset of 20 spiders using DNeasy Blood & Tissue Kit (Qiagen) following the manufacturer’s protocol. We performed diagnostic PCR for *Rickettsiella*, and two other symbiont genera (*Tisiphia* and *Wolbachia*) known to be present in some *M. fradeorum* using previously published protocols (wsp primers for *Wolbachia*, Baldo et al. 2006; Ricklong primers for *Tisiphia*, Curry et al. 2015; RLA primers for *Rickettsiella*, Rosenwald et al. 2020). All tested spiders were positive or negative for *Rickettsiella* as expected, and all spiders were negative for the other symbionts.

To evaluate the hatch rate, we dissected each eggmass after spiderlings emerged, and counted the remaining unhatched eggs as in Rosenwald et al. (2020). Total hatched and unhatched eggs per female were used for a series of logistical regression contrasts, using Williams’ correction for overdispersion (Arc v 1.06; Williams 1982). In the CI induction experiment, we conducted three statistical contrasts. First, to ensure CI was in evidence in the experiment, we compared hatch rates in expected CI treatments (warm and cool R-infected males mated to uninfected females) to control matings between uninfected males and uninfected females. Second, to ensure that neither temperature nor *Rickettsiella* infection negatively affected male fertility, we then contrasted hatch rates when these same males were mated to cool-reared *R*-infected females, relative to uninfected males mated to R-infected females. Finally, we directly contrasted the hatch rates of the warm and cool CI treatments to one another, to evaluate the effect of rearing temperature on CI induction. In the CI rescue experiment we likewise conducted three statistical contrasts. First, to ensure rearing temperature didn’t directly affect hatch rate, we contrasted hatch rates from warm versus cool R-infected females when mated to uninfected males. Second, to ensure CI was in evidence in the experiment, we contrasted cool R-infected females mated to cool R-infected males (cool rescue treatment), relative to cool uninfected females mated to cool R-infected males (CI control). Finally, we directly contrasted hatch rates of the warm and cool CI rescue treatments to one another, to evaluate the effect of rearing temperature on CI rescue.

We evaluated symbiont titer via qPCR for a subset of 10-15 R-infected spiders from each sex and temperature treatment. For the sake of consistency, all evaluated spiders originated from the same experiment, the CI rescue experiment. We placed specimens into 50% PBS (EtOH: PBS) for 30 min, followed by 75% PBS (EtOH: PBS) for 30 min, and finally kept in 100% PBS overnight. We crushed these specimens in cryogenic vials containing solid-glass beads using FastPrep® Tissue Homogenizer (MP Biomedicals), then extracted DNA using DNeasy Blood & Tissue Kit (Qiagen) following the manufacturer’s protocol. We designed short primers specific to the *Rickettsiella recA* gene and *M. fradeorum 18S rRNA* gene using Geneious Prime Version 2021.2 (Table S1). Primer specificity was tested *in-silico* and using diagnostic PCR (Table S2). We cloned these short specific sequences using a vector from CloneJET™ PCR Cloning Kit and JM109 Mix & Go Competent Cells. Serial dilutions with serial factor 1:10 for the symbiont gene and 1:3 for the spider gene were used for a standard calibration curve. qPCR was performed using the Step One Plus real-time PCR system (Applied Biosystems, CA, USA) and Fast SYBR green master mix (Thermo Fisher Scientific, MA, USA). For each 20 μL reaction mixture, we used 30 ng of gDNA, and each sample was measured in three technical triplicates. Cycling conditions are listed in Table S2. We used output Ct, Intercept, and Slope values to calculate the amount (ng) of the symbiont’s gene and the spider’s gene from each sample 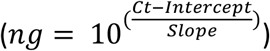. We then used these weights to calculate the gene copy number (gcn) 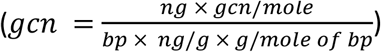, followed by data normalization 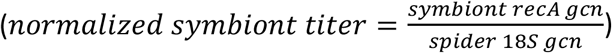.We log-transformed the normalized titer values to reduce heteroscedasticity, and compared means using a general linear model using Type III sums of squares in IBM SPSS v 28.0., the normalized titer values to reduce heteroscedasticity, with sex, temperature, and their interaction included as fixed factors.

## RESULTS

When we reared R-infected spiders at either 26°C or 20°C and then evaluated the strength of CI induction (Figure 1A), we found that both warm and cool R-infected males caused reduced hatch rates when mated to uninfected females, relative to control matings between uninfected males and females (ΔDeviance = 58.95, d.f. = 2, P<0.001). When these same R-infected males were mated with R-infected females, hatch rates were nearly perfect and did not differ from control crosses between uninfected males and R-infected females (ΔDeviance = 2.87, d.f. = 2, P=0.238), indicating that neither rearing temperature nor *Rickettsiella* infection intrinsically decreased fertility of males. When compared directly, the hatch rate of incompatible crosses between uninfected females and warm males was twice as high (0.687±0.052) as with cool males (0.348±0.046; ΔDeviance = 21.30, d.f. = 1, P<0.001), indicating that warmer rearing temperatures weakened CI induction.

**Figure 1.**
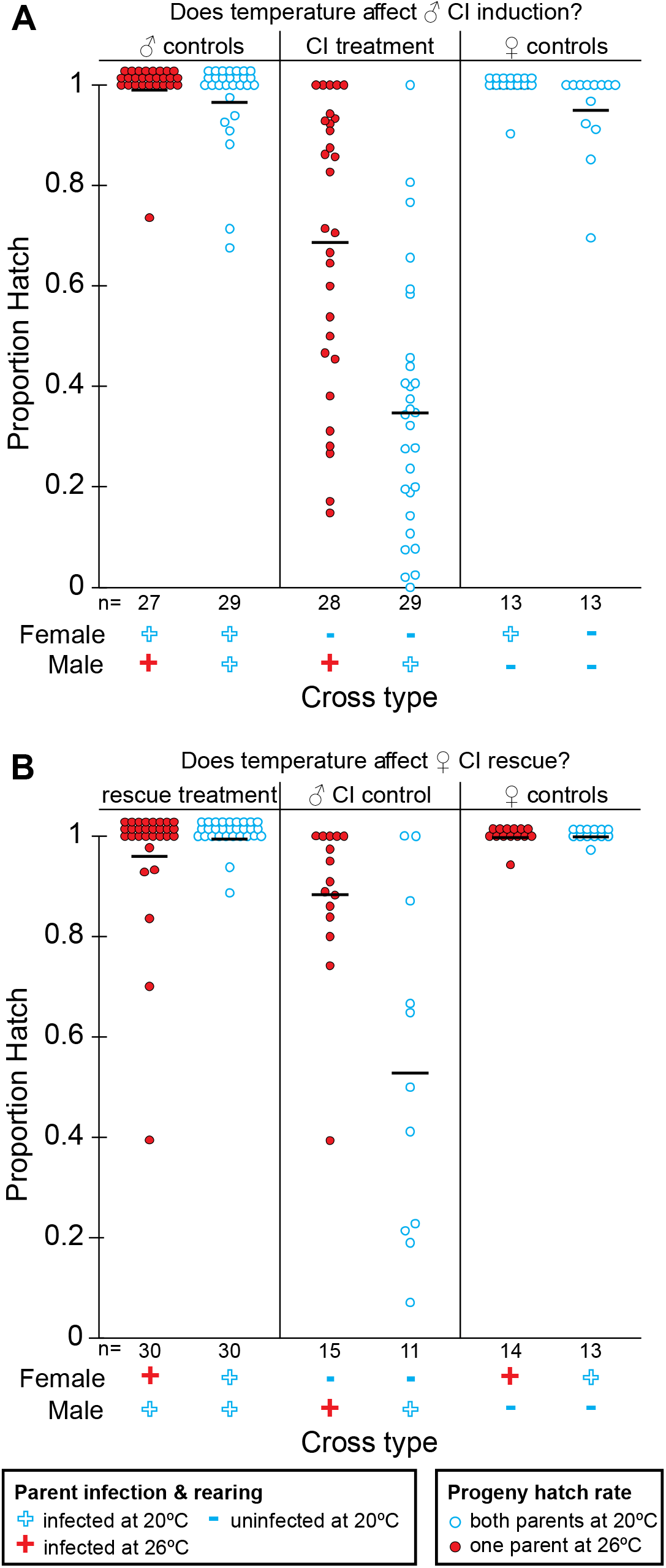
Proportion hatch of eggs from crosses between *Rickettsiella*-infected *Mermessus fradeorum* spiders that were individually reared at 26°C or 20°C, and subsequently mated to either R-infected or uninfected spiders reared at 20°C as indicated on the x-axes. Realized sample sizes per cross (excluding failed matings) are indicated below each column. Crosses to evaluate male CI induction (Panel A) showed a significantly higher hatch rate (weaker CI) for offspring of warm-reared than cool-reared infected males when mated to uninfected females. All control crosses had high hatch rates. Crosses to evaluate female CI rescue (Panel B) showed marginally lower hatch rate for warm-reared than cool-reared infected females mated to infected males, but high hatch rates overall indicate temperature had relatively little effect on CI rescue. Incompatible crosses between infected males and uninfected females were included as controls to aid comparison across the two experiments and validate that infected males used within this experiment were capable of inducing CI.

In the CI rescue experiment (Figure 1B), hatch rates were high and equivalent for warm and cool R-infected females mated to uninfected males (ΔDeviance=0.20, d.f.=1, P=0.654), indicating that rearing temperatures did not directly affect hatch rate. Cool R-infected females also exhibited nearly perfect hatch rates when mated to cool R-infected males, much higher than cool uninfected females mated to cool R-infected males (ΔDeviance=84.49, d.f.=1, P<0.001). This control contrast establishes both that the R-infected males in this experiment induced CI, and that cool R-infected females were capable of rescue. When we contrasted rescue between the two female rearing temperatures, we found that maternal rearing temperature had a marginally significant effect on hatch rate (ΔDeviance = 3.825, d.f. =1, P=0.0505), suggesting that CI rescue ability can be weakened in at least some R-infected females. Nevertheless, most females in these rescue crosses exhibited perfect hatch rates, indicating that the CI rescue ability of *Rickettsiella* usually remains intact regardless of rearing temperature.

When we evaluated bacterial titer in a subset of warm-reared versus cool-reared spiders (Figure 2), we found an interactive effect between sex and temperature (Sex F_1,44_ =0.329, P=0.529, Temperature F_1,44_ =6.026, P=0.018, Sex × Temp interaction F_1,44_ =5.22, P=0.027). Titer was lower in the warm treatments, but this lower titer was associated with females more than males, even though temperature effects on symbiont phenotype were more evident in males (CI induction) than females (CI rescue).

**Figure 2.**
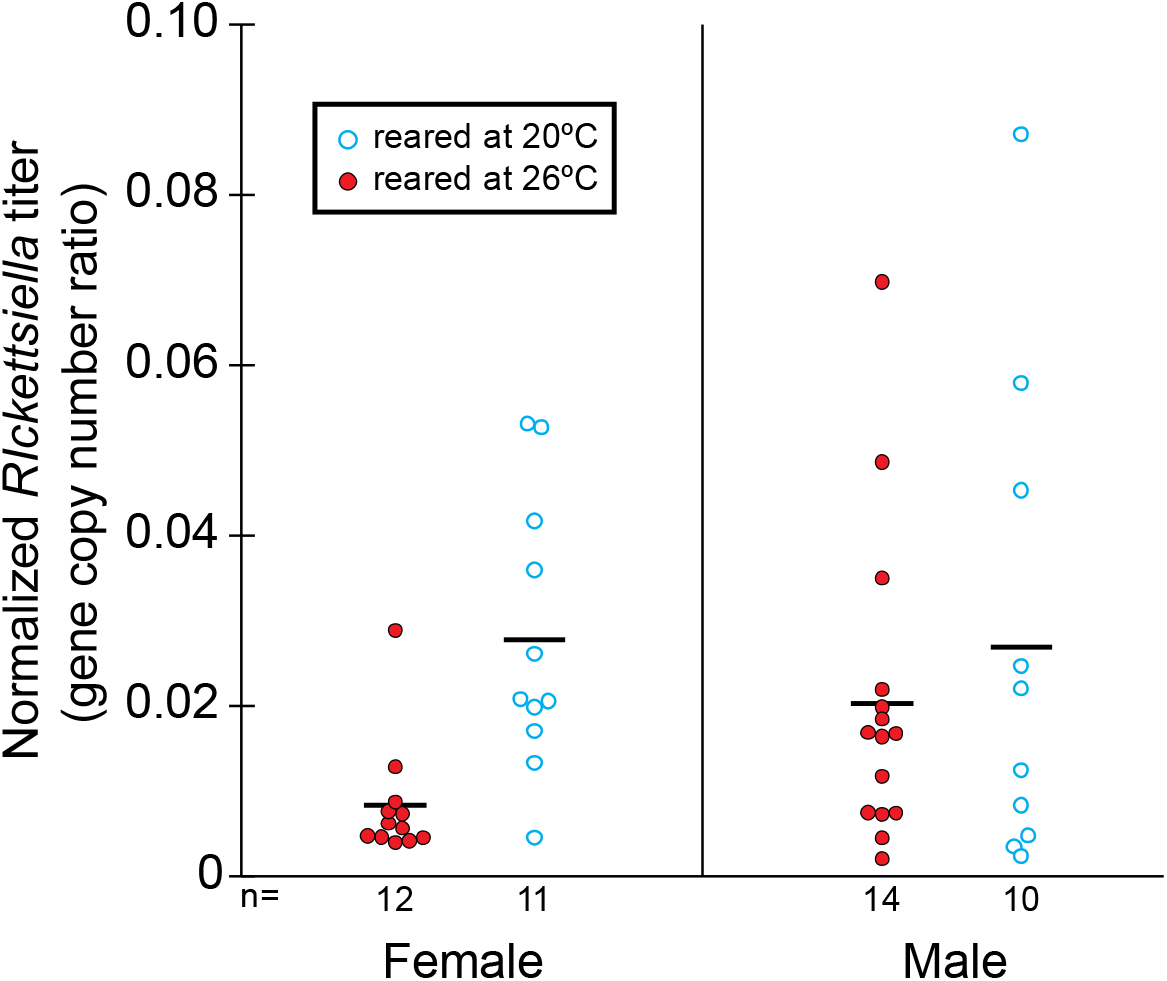
*Rickettsiella* titer in a subset of warm (26°C) and cool (20°C) reared spiders from the CI rescue experiment. Log transformed titer (normalized against spider 18S quantity) was significantly affected by the interaction between temperature and sex, primarily driven by lower titer in warm reared females.

## DISCUSSION

Warmer rearing temperatures strongly decreased CI induction by *Rickettsiella* in male *M. fradeorum* spiders, but only mildly decreased CI rescue in female spiders. Thermal sensitivity for *Rickettsiella*-induced phenotypes is consistent with results for a different strain of *Rickettsiella* in an aphid host, which exhibits a different set of host phenotypic effects that don’t include CI (Gu et al. 2023). In the present study, CI strength overall was notably weaker than in prior studies by our group (Rosenwald et al. 2020), which were conducted at room temperature, but in a laboratory that was cool throughout the duration of the experiments (J. White, pers. obs.). Highly variable hatch rates across all CI crosses (Fig 1A), suggest that other factors may additionally be driving CI strength in this system (e.g., Shropshire et al. 2021). More generally, bacterial symbionts often show reduced phenotypic penetrance at warmer temperatures (Martins et al. 2023), although the phenomenon is not universal, and instances where symbiont-induced phenotypes increase with warmer temperatures also exist (e.g., Corbin et al. 2021).

Warmer rearing temperature also decreased *Rickettsiella* titer, particularly in female spiders, which also tended to have lower titer than males. The latter was surprising, because gravid females are usually expected to carry higher loads of maternally transmitted symbionts than males (e.g., Noda et al., 2001). However, it is also possible that relatively high loads of bacteria would be allocated to eggs, and these post-ovipositional females, which had laid 3 eggmasses prior to preservation and extraction, may have exhibited substantially reduced bacterial titer relative to their pre-ovipositional state. It is also worth noting that our standard laboratory protocol calls for imposing several days of food deprivation to clear gut contents (and associated prey symbionts) before spider specimen preservation. This protocol likely diminished the utility of the qPCR assay, by temporally disassociating the titer of the preserved specimen from its titer at biologically relevant timepoints (fertilization, egg deposition). Despite these caveats, we nevertheless did detect a signal of temperature treatment on bacterial titer, suggesting that stronger CI penetrance may be associated with higher bacterial titer. We will be following up with assays more specifically designed to probe the mechanism and timing of CI modification and rescue in this novel host/symbiont system. Associations between temperature and symbiont titers have previously been demonstrated with other endosymbionts (Breeuwer and Warren 1993, Lopez-Madrigal and Duarte 2019, Ross et al. 2020), and thus our results further support the potential links between temperature, titer, and phenotypic outcome.

Our study demonstrates the thermal sensitivity of CI in a novel arthropod/symbiont system, implying that temperature may be modulating host/symbiont interactions quite broadly in nature (e.g., Kreisner et al. 2016, Hague et al. 2022). However, much additional study will be needed before we can predict how temperature affects CI dynamics in the field. For *M. fradeorum*, we selected only two relatively moderate temperatures to test, which the spider likely experiences routinely in the field. We also kept each temperature constant, because we do not yet know the mechanism or timing of *Rickettsiella*-induced CI modification and rescue. Future studies will explore variations of timing, duration, and extent of thermal exposures on CI in this system, which will inform our understanding of spider/symbiont biogeography for this widespread introduced species within the context of global climate change.

More broadly, environmental effects on symbiont phenotypic penetrance may have important implications in both managed and natural ecosystems. *Wolbachia* is already in use in vector management in several areas of the globe, with mixed results (Hoffmann et al. 2011, Dos Santos et al. 2022, Hoffmann et al. 2024). Symbiont thermal tolerance is one factor that appears to limit the success of this technique in some regions (Ross et al. 2020, 2023). As the use of symbionts to control pests and vectors comes under consideration in a range of other systems (e.g., Gong et al. 2020, Mateos et al. 2020), it becomes critical to characterize the environmental constraints that might influence efficacy. Even in less-managed systems, the thermal optima of symbionts might play a critical role in determining the distribution of pests and the ability of natural enemies to control them (Zhang et al. 2019, Hague et al. 2022). Our study contributes to a growing body of literature that suggests environmental contingency may be the rule for interactions between hosts and reproductive manipulators, not the exception.

## ACKNOWLEDGEMENTS

We thank L. Rosenwald, K. Butler, and E. Williams for assistance in the lab, and M. Doremus and three anonymous reviewers for helpful comments on previous versions of this manuscript.

