## Supplemental Document 1 for "Warm temperature inhibits cytoplasmic incompatibility induced by endosymbiotic *Rickettsiella* in a spider host"

**Supplemental Document 1: PCR primers and cycling conditions**

Table S1. Primers used for quantitative PCR.

| Target | Target gene | Primer name | Primer sequence 5' to 3' | Expected product size (bases) | Annealing temp |
|---|---|---|---|---|---|
| *Rickettsiella* | *recA* | recA_345F<br>recA_510F | ACAACCTGATACTGGCGAGC<br>CGACATTAATCGCGCTTGCA | 166 | 60 $^{o}$C |
| *M. fradeorum* | *18S* | Mfra18S_876F<br>Mfra18S_1007R | ACAACTGCCCGTTCTGAACA<br>GCACCATTGTTCAGGCCTTG | 132 | 60 $^{o}$C |

Table S2. PCR and qPCR cycling conditions

| | Cycling program | | Mixture per reaction | |
|---|---|---|---|---|
| **PCR** | Initial denaturation | 95°C for 3 min | GoTaq® Green Master Mix | 10 µL |
| | Denaturation | 95°C for 30 s | 10µM Forward Primer | 1 µL |
| | Annealing | 60°C for 24 s | 10µM Reverse Primer | 1µL |
| | Extension | 72°C for 1 min | DNA Template | 30ng |
| | Final extension | 72 $^{o}$C for 10 min | Nuclease-Free Water | To 20 µL |
| **qPCR** | Holding Step 1 | 95°C for 20 s | SYBR™ Green Master Mix | 10 µL |
| | Cycling Step 1 | 95°C for 3 s | 10µM Forward Primer | 0.8 µL |
| | Cycling Step 2 | 60°C for 30 s | 10µM Reverse Primer | 0.8 µL |
| | Melt curve Step 1 | 72°C for 15 s | DNA Template | 30ng |
| | Melt Curve Step 2 | 72 $^{o}$C for 60 s | Nuclease-Free Water | To 20 µL |
| | Melt Curve Step 3 | 95 $^{o}$C for 15 s | | |
